# KuLGaP: A Selective Measure for Assessing Therapy Response in Patient-Derived Xenografts

**DOI:** 10.1101/2020.09.08.287573

**Authors:** Janosch Ortmann, Ladislav Rampášek, Elijah Tai, Arvind Singh Mer, Ruoshi Shi, Erin L Stewart, Celine Mascaux, Aline Fares, Nhu-An Pham, Gangesh Beri, Christopher Eeles, Denis Tkachuk, Chantal Ho, Shingo Sakashita, Jessica Weiss, Xiaoqian Jiang, Geoffrey Liu, David W. Cescon, Catherine O’Brien, Sheng Guo, Ming-Sound Tsao, Benjamin Haibe-Kains, Anna Goldenberg

## Abstract

Quantifying response to drug treatment in mouse models of human cancer is important for treatment development and assignment, and yet remains a challenging task. A preferred measure to quantify this response should take into account as much of the experimental data as possible, i.e. both tumor size over time and the variation among replicates. We propose a theoretically grounded measure, KuLGaP, to compute the difference between the treatment and control arms. KuLGaP is more selective than currently existing measures, reduces the risk of false positive calls and improves translation of the lab results to clinical practice.

---

Despite tremendous advances in pharmaceutical research, many cancer patients do not respond to the first line of therapy. In oncology, researchers rely on preclinical models to investigate drug response to assess whether a drug works against a given cancer type. Of the available preclinical models, *in vivo* models tend to capture response to the drugs more faithfully than *in vitro* models ^1^. A standard readout from *in vivo* models is the size of the tumor growth over time across multiple experimental replicates compared to a set of untreated controls. As with most biological systems,, tumor growth can vary within and between host mice creating substantial variance among biological replicates. Determining whether the *in vivo* model is actually responsive to the given drug from the set of biological experiments is thus a complex task. It is essential, however, to make an accurate determination as it has a direct impact on translation to the clinic.

Many measures have been proposed to quantify response to a treatment for *in vivo* models^1–3^. Commonly used measures include mRECIST, area under the curve (AUC) ^2^, angle of response (Angle) ^2^ and the tumor growth inhibition (TGI) ^3–5^. Depending on which measure a researcher selects, the assessment of response may yield different, often opposite, conclusions, as none of the existing measures take full advantage of the data collected across replicates. For example, mRECIST ^1^ is easy to compute but does not take controls into account and is thus unable to distinguish true disease control (stable disease) from a naturally slow-growing tumor. The angle of response and TGI only take into account the last measurement rather than the full trajectory of treatment. All of AUC, angle of response and TGI measures ignore variance in the replicate experiments, depending only on the mean across replicates. These limitations often lead to an over-optimistic assessment of response, therefore failing to faithfully recapitulate clinical observations as can be seen from our experiments.

In this work, we show how multiple sources of variation lead to erroneous response calls and propose a new response measure, KuLGaP (based on **Kul**lback-**L**eibler divergence between **Ga**ussian **P**rocesses), which accounts for both experimental controls and variation among replicates. We test and compare KuLGaP to four widely-used response measures using 329 patient-derived xenograft (PDX) models and show that our measure leads to a more conservative response call rate with better concordance to clinical response in patients in a small trial cohort. Indeed, in a cohort of 13 patients and their xenografts, the mean time to relapse (MTR) in patients we identify as responders from their PDX according to our KuLGaP measure is at least twice as high as MTR for responders identified by other measures. Our KuLGaP statistic bridges the gap between the potential of PDX data and its clinical application by utilizing the full extent of the experimental data. Last but not least, we show that the robustness of KuLGaP allows for experimental designs with fewer animal replicates without significant loss in the accuracy of response quantification.

## Results

To illustrate the benefits and pitfalls of various measures assessing the drug response in PDX, we collected tumor growth curves from 329 PDX models (Supplementary Table 1). In the following analysis, schematically captured in Figure 1, we evaluate our new KuLGaP growth response measure compared to other commonly used response measures: mRECIST ^1^, area under the curve (AUC)^6^, angle of response (Angle) ^2^, and tumor growth inhibition (TGI) ^3–5^ and show that by using experimental data more comprehensively, we achieve a more selective measure that better corresponds to patient treatment response.

**Table 1.** Patient stratification by corresponding PDX response. Mean time to relapse (in years) in the group of responders and non-responders according to each measure. The number of patients is indicated in parentheses.

| Measure | Mean time to relapse in responders | Mean time to relapse in non-responders | Difference (years) |
| --- | --- | --- | --- |
| KuLGaP | 4.13 (3) | 1.02 (10) | 3.11 |
| mRECIST | 2.62 (7) | 0.97 (6) | 1.65 |
| Angle | 2.14 (10) | 1.53 (3) | 0.61 |
| AUC | 2.02 (11) | 0.32 (2) | 1.70 |
| TGI | 2.22 (7) | 1.25 (6) | 0.97 |

**Figure 1:**
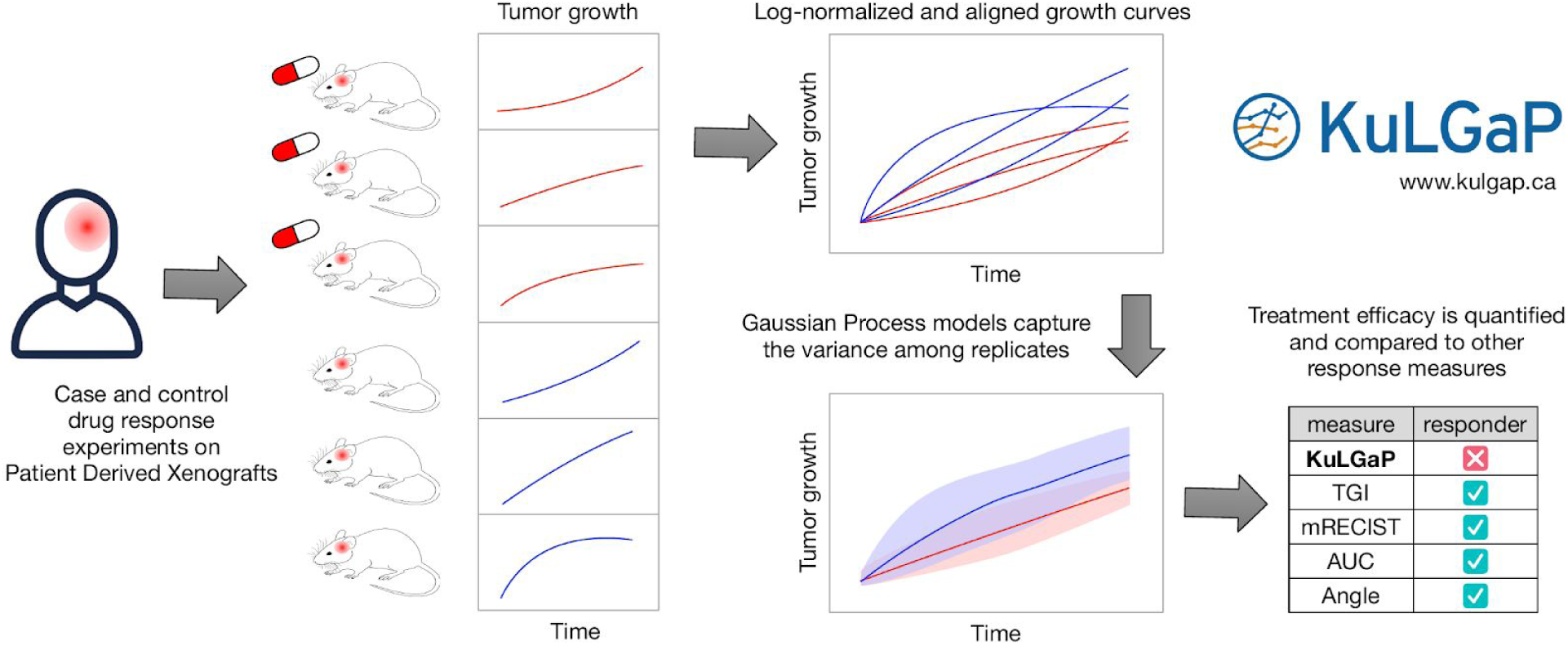
The KuLGaP pipeline overview. The human tumor is implanted in a set of mouse replicates (patient derived xenografts; PDX) of which some are treated with a given drug (cases; in red) and some are not (controls; in blue). In addition to KuLGaP we also evaluate four different commonly used measures to assess response status in PDXs. Since none of the existing approaches fully account for both variation among replicates and the baseline growth of controls, they tend to over-optimistically indicate that a PDX model was a responder. We designed KuLGaP to address these shortcomings: taking the variance among replicates into account in a statistically rigorous manner leads to a more accurate response categorization.

### KuLGaP, a new measure for in vivo therapy response

There are two steps to computing KuLGaP. First, two Gaussian Process (GP) models ^7^ are fitted to the PDX tumor growth curves, one for treated PDXs and another for controls. Second, we compute the distance between these two GP models using Kullback-Leibler (KL) divergence ^8^. Schematically this process is captured in Figure 1 as well as Supplementary Figure 1. The benefit of using the GP models is that they not only model the covariance of measurements across time ^7^, but they also model the variance within a group of replicates over time. This, as we show in the paper, is necessary for robust identification of responders. KL divergence that we use to compare treated replicates and controls, is often used in machine learning and mathematical fields to measure the difference between two distributions, as KL has strong theoretical foundation in information theory ^9^ and can be quickly computed for many distributions ^10^, including the Gaussian distribution.

We assess the significance of the distance between a treatment and control arms by computing an empirical null distribution of distances between all pairs of controls in our dataset. Using this empirical distribution, we compute the significance (p-value) of treatment response for each PDX model. Models with a p-value less than 0.05 are considered to have a statistically significant distance and are classified as responders. The computation of KuLGaP is illustrated in Supplementary Figure 1 and is described in depth in Online Methods.

### Comparison of measures for therapy response

We performed a comparative analysis of KuLGaP with mRECIST, AUC, Angle and TGI. For each pair of measures, we computed the level of agreement between them as the percentage of experiments on which both measures give the same classification (responder or non-responder) (Figure 2a), the false discovery rate of each measure with respect to each other (Figure 2b) and related the number of compared measures that classified an experiment as a responder to the KuLGaP statistic (Figure 2c).

**Figure 2:**
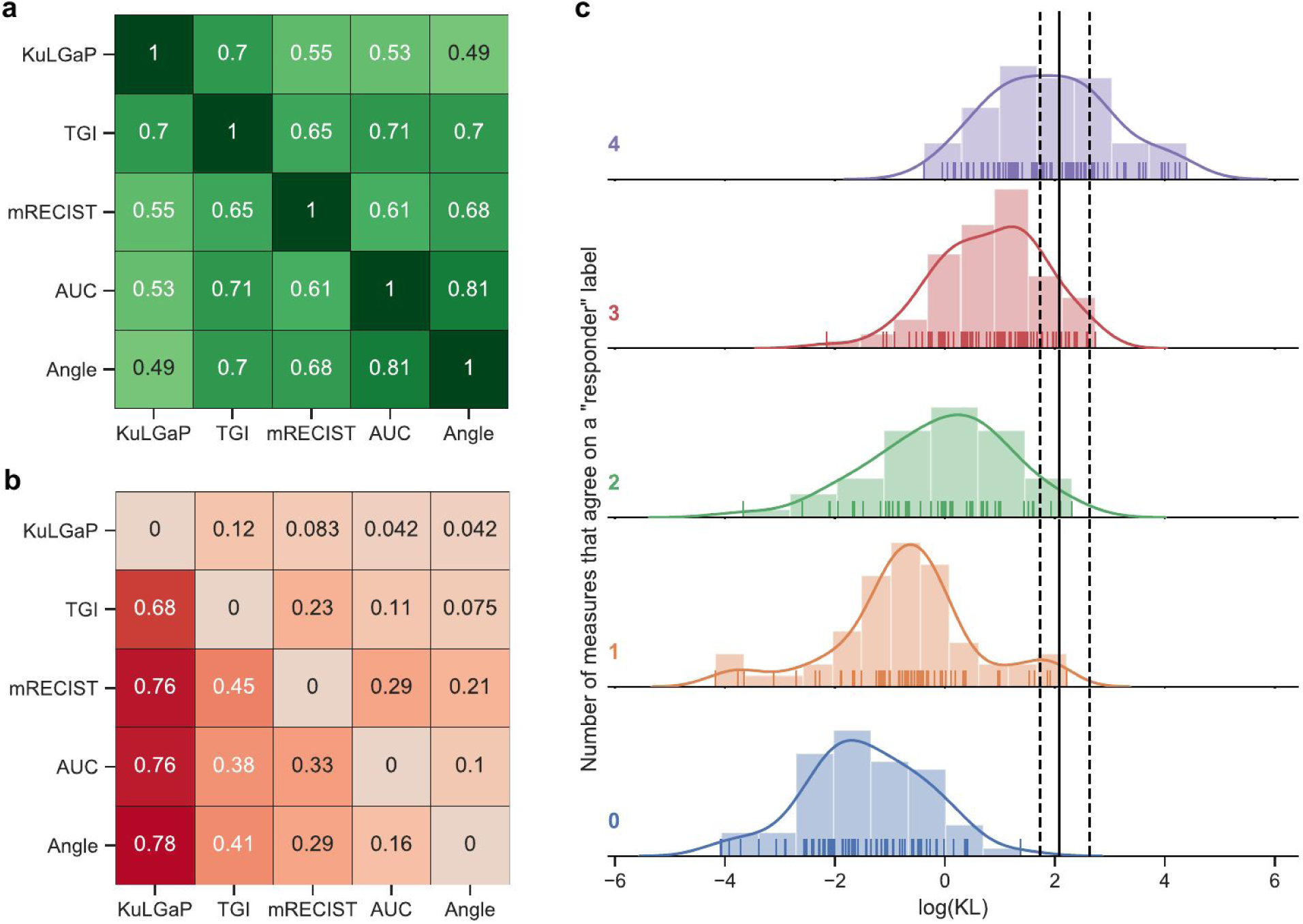
Comparison of classifications according to all 5 response measures. **a**, Heatmap of agreement (fraction of experiments where two measures agree) between the different measures across all models. **b**, Proportion of responders according to the measure in the row that were not considered responders by the measure listed in the column. **c**, Each row shows a histogram (distribution) of KuLGaP KL-values across a group of experiments for which (top to bottom): i) all baseline measures (TGI, mRECIST, AUC, Angle) agreed on responder classification for each experiment in this group (top, purple); ii) three out of four baseline measures agreed on responder classification for experiments in this group (red); iii-v) 2–0 (green – blue) baseline measures agreed on a responder classification, respectively. The solid vertical line indicates the KuLGaP’s threshold for significance (calling an experiment a responder) at the 0.05 level, while the dashed lines indicate the 0.1 and 0.001 thresholds, respectively. All experiments to the right of the vertical line are responders according to KuLGaP and all experiments to the left are non-responders according to KuLGaP.

Out of the 329 experiments (with a total of 1437 treatment and 1946 control arms), KuLGaP classified 48 as responders (14.6%), compared to 133 (40.4%) for TGI, 187 (56.8%) for mRECIST, 186 (56.5%) for AUC and 211 (64.1%) for Angle. Briefly, TGI and Angle depend on the ratio of the difference between the first and last growth measurement of treatment and control; AUC calculates the cumulative difference at each measurement point between control and treatment groups; mRECIST categorises observation of growth in the treatment arm into Complete Response (mCR), partial response (mPR), stable disease (mSD) and progressive disease (mPD). Following established practice ^11–13^, we consider all PDX with a TGI value of more than 0.6 to be responders. Please see Online Methods for a detailed definition of each measure. The measures that give the most similar results are AUC and Angle. Our KuLGaP measure yields results that are most similar to TGI: the two classifications agree on responders and non-responders in 70% of all cases. Overall, KuLGaP is more conservative than all other response measures (Figure 2b), indicated by fewer responders called as compared to other methods. For example, all but 4 experiments that are classified as responders by KuLGaP (92%) are also responders according to mRECIST, whereas only 31 out of 147 mRECIST-responders are also responders according to KuLGaP. Similarly, all but two of the KuLGaP responders are called responders by Angle and AUC, but each of these measures call many KuLGaP-nonresponders as responders (141 for AUC, 165 for Angle). Figure 2c shows that there is significant disagreement between the different measures. However, KuLGaP captures the majority of experiments for which there is a consensus among the other measures.

Further, we compared the continuous measures underlying TGI and KL classifications. We found that the Spearman rank correlation coefficient between TGI and the logarithm of the Kullback-Leibler divergence is 0.69. However, TGI suffers from some of the drawbacks. We identified two main factors contributing to the observed inconsistent calls by the various measures: incorporation of the control group information and variance across replicates. We use examples from our data in order to illustrate the importance of these factors next.

### Importance of the control group

We found that the information contained in the control replicates is crucial for an accurate response classification. A downside of the mRECIST classification is that it does not consider the control group but makes a classification based on the treatment group alone. The mRECIST criterion rates each treatment replicate as either complete response (mCR), partial response (mPR), stable disease (mSD) or progressive disease (mPD), and then classifies the experiment by a majority vote of all the replicates where all but mPD ratings are considered a responder ^1^. In Supplementary Figure 2, we show an extreme example of how over-optimistic mRECIST would be if mSD were also considered a responder.

Consider the following two NSCLC PDX models ^14^, Model 1 (Figure 3a-c) treated with afatinib and Model 2 (Figure 3d-f) treated with erlotinib, respectively. The mRECIST framework reports stable disease (mSD) for all replicates of both models, resulting in a “responder” call for both models, while KuLGaP called a significant response for Model 2, but not Model 1. Because mRECIST does not take into account the control group, it missed the fact that the cancer in the untreated arm grows as fast as in the treated arm in Model 1, but not in Model 2. Therefore the mRECIST classification is the same for both models, despite the clear differences. Both the AUC and Angle classifications agree with the KuLGaP classification in both cases, whereas TGI classifies both models as non-responders. Because TGI does not consider the length of time for which the treated sample is not growing, it fails to detect the tumor arrest in Model 2. In Model 2, the treatment arm of this model took approximately 50 days longer to reach the maximum tumor size of its control replicates, and this effect was detected by our KuLGaP approach.

**Figure 3:**
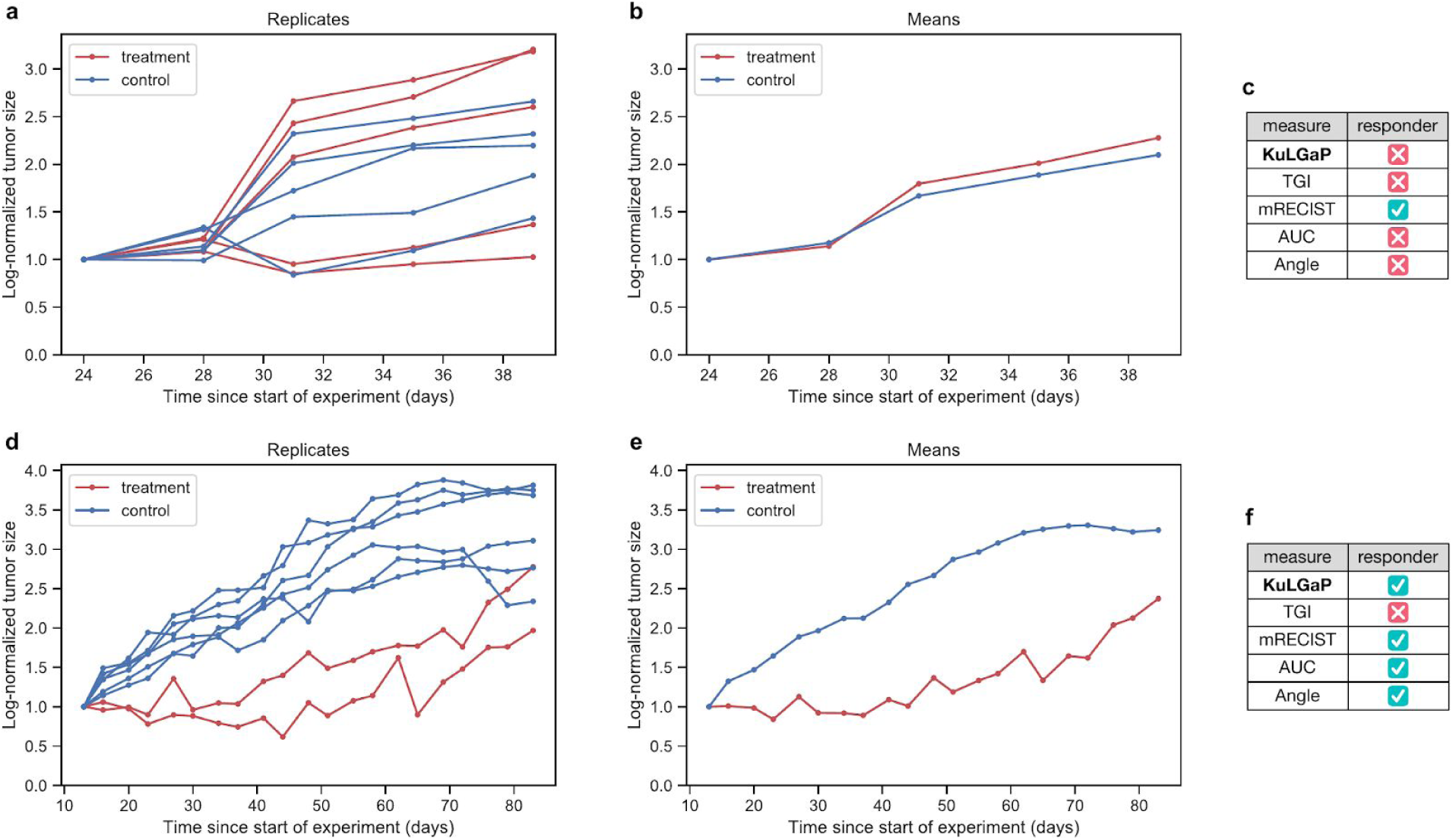
Importance of the control group. **a**, Log-normalized growth curves (afatinib treatment arm in red, control arm in blue) of a NSCLC PDX model (Model 1) with five replicates in each arm ^14^. **b**, Means across treatment and control replicates of Model 1 from panel **a. c**, Classification of Model 1 response to the treatment. **d**, Log-normalized growth curves of another NSCLC PDX model (Model 2) with two erlotinib treatment replicates and six controls ^14^. **e**,**f**, Analogous to panels **b** and **c**, respectively, but for Model 2. The mRECIST measure identifies both models as responders, particularly as stable disease (mSD); KulGaP identifies Model 1 as a non-responder and Model 2 as a responder.

For Model 1 (Figure 3a-c) we observed a particularly over-optimistic responder call by mRECIST; another such example is shown in Supplementary Figure 2. An intuitive way to alter the mRECIST classification to be more conservative is to consider only the mCR and mPR ratings as a positive response. However this leads to considerable loss of sensitivity, as demonstrated in Supplementary Figure 3. The simple alteration cannot fix a fundamental mRECIST flaw.

Furthermore, in Supplementary Figure 4a-c we show a colorectal cancer PDX with 8 control and 8 treatment replicates treated with evofosfamide. All measures apart from KuLGaP classified this model as a responder. The mRECIST measure fails to take into account the fact that the treatment and control groups grow at a similar pace, whereas Angle and AUC only consider the last day of measurement and therefore miss the greater similarity of the treatment and control growth curves throughout the experiment. We provide an additional example supporting our claims in Supplementary Figure 4d-f.

### Accounting for variance among replicates is important

Accounting for the variance among replicates leads to greater selectivity in declaring a response. An illustration of this scenario is given by the breast cancer PDX experiment with 15 paclitaxel treated and 12 control replicates shown in Figure 4a-c. While there is a substantial difference in the means between the control and treatment groups (Figure 4b), there is also significant variance among replicates in each group (Figure 4a). TGI, just like mRECIST, AUC and Angle measures, classifies this model as a responder. Our KuLGaP takes into account the variance among replicates and shows that the variance within control and treatment arms is big enough to remove the significance of the mean difference, thus classifying this model as a non-responder.

**Figure 4:**
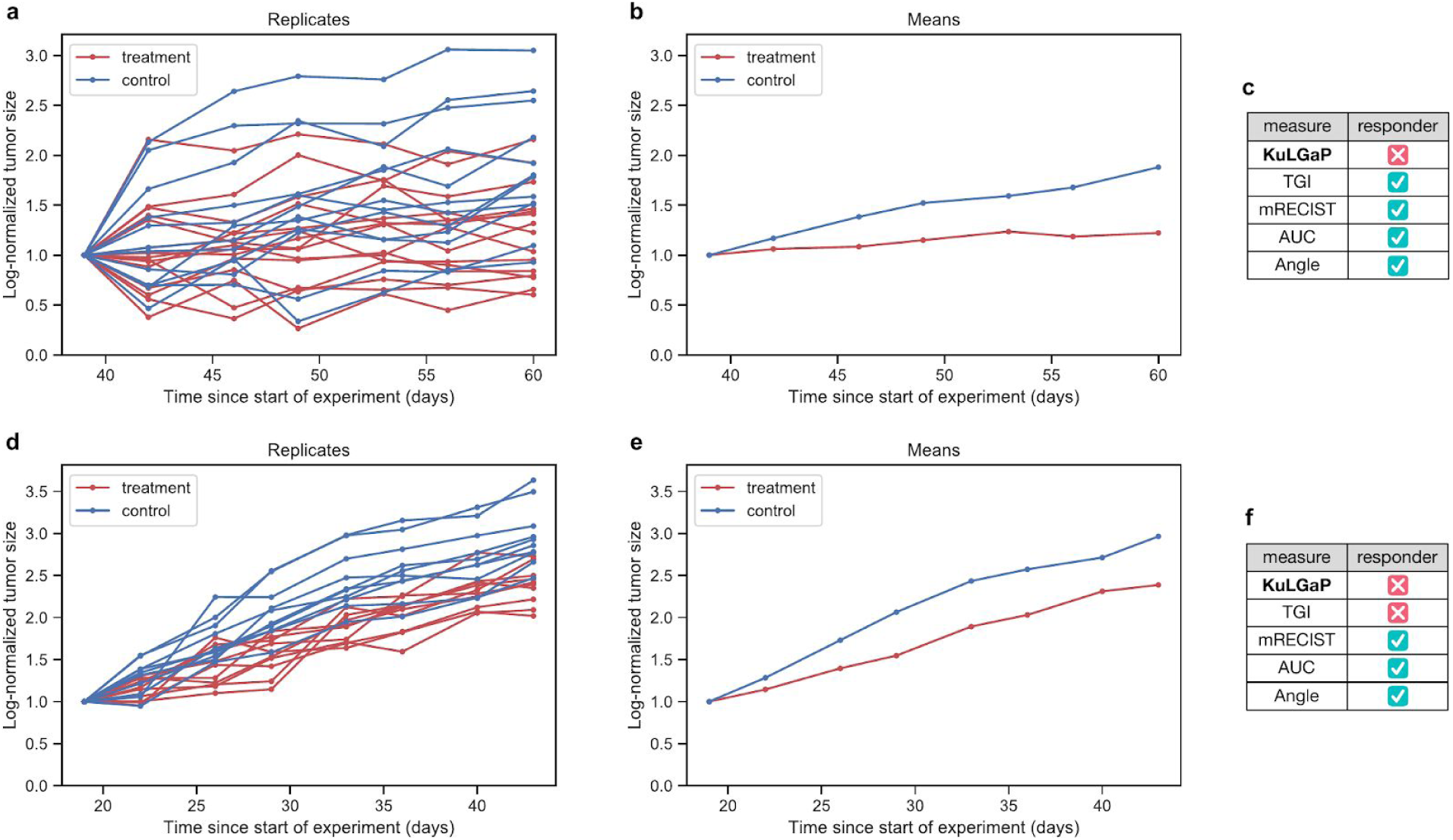
Importance of accounting for variance. **a**, Log-normalized tumor growth curves of a breast cancer PDX model^15^ treated with paclitaxel; 15 treatment (in red) and 12 control (in blue) replicates. **d**, Log-normalized growth curves of a NSCLC PDX model with 10 replicates treated with dacomitinib. **b**,**e**, Mean treatment and control arm growth curves for each model (**a**,**d**), respectively. **c**,**f**, Computed response classifications by all compared response measures for each model (**a**,**d**), respectively.

Next, consider the following experiment, where 10 replicates of an NSCLC PDX model were treated with dacomitinib (Figure 4d-f). The Angle and AUC measures, which do not take into account variance, identify this PDX model as a responder. Our KuLGaP measure picks up on the fact that the variance among replicates in the treatment and control groups is larger than the mean difference between the two groups, and therefore declares the experiment a non-responder. In other words, incorporation of variance leads to greater selectivity in declaring response. The TGI measure concurs with the KuLGaP assessment of a non-responder. The mRECIST classification (which does not consider the control group) is stable disease (mSD), and thus the model is erroneously considered responsive.

An additional example, provided in Supplementary Figure 5, shows an experiment where even a large difference between the mean growth of the treatment and control arms can be deceptive. Upon closer inspection of the individual replicates it is clear that any difference in the mean behaviour is dwarfed by the large variance, leading to a false positive call by all measures but KuLGaP.

### Implications of not considering multiple replicates in the study design

The experimental design of xenograft experiments usually requires the researcher to collect responses from multiple replicates of the model treated with the drug comparing them to those that are treatment-naive (controls). Since, particularly using PDX, these experiments are laborious, a 1×1×1 experimental design was proposed ^1^, where only a single replicate is used per drug and model. By testing and publishing a dataset on 1000 PDXs, the NIBR PDXE study greatly contributed to research in this area. Unfortunately, this experimental design has its limitations. In this setup the researchers are able to gain insight into the population-level response for a given drug. However, this design is not sufficient to draw conclusions for an individual patient (that the PDX was derived from) level due to the absence of the variability that can only be derived from the replicates of the same PDX ^16^.

The lack of accounting for the variance in the 1×1×1 design is particularly detrimental for the mRECIST classification used in the study^1^. Indeed, it is common for different replicates to have different mRECIST classifications. An extreme case is given by an experiment with five treatment replicates (see Supplementary Figure 6). Two of the five replicates are classified as mPR, two as mSD and one as mPD. Depending on the one randomly chosen replicate in the *n=1* design, the classifications would have been different. This scenario is common since the mRECIST classification is often decided early on in the experiment, when the tumors are smaller, and therefore more susceptible to measurement errors and noise. In our dataset, we find that fewer than 30% (97 out of 329) of the models have the same mRECIST classification across replicates. Almost 60% (197 out of 329) of the models have two different mRECIST classifications, of which 39 models (11% of the total) have mRECIST classifications that are not adjacent (such as mCR and mSD). In 10% (32 out of 329) of the models, treatment replicates are assigned three different mRECIST classifications. The resulting number is staggering: almost half (160 out of 329) of the models have a majority decision that is supported by fewer than 75% of replicates of that model. Consequently, we postulate that the NIBR PDXE study using the 1×1×1 design with mRECIST criterion is likely to be unreliable for personalized treatment prediction in many clinical scenarios.

### Assessing a study design with fewer replicates

There is a significant downside to having only a single replicate per experiment. However, a large number of replicates increases the cost, and the use of research animals. We performed a further experiment to see whether a smaller number of replicates would achieve reliable results. For each experiment, we randomly sampled without replacement three treatment and three control replicates and computed the KuLGaP, mRECIST, Angle, AUC and TGI classifications based on this sub-sample. This was repeated three times. Thus, for each model, we obtain 3 sets of experiments with 3 replicates each. By comparing the responses using only 3 replicates to those obtained using the full set of replicates we were able to estimate how robust each response measure is to a reduced number of replicates. We found that KuLGaP and TGI measures are particularly robust to this form of subsampling, reaching agreements of 95.9% and 94.1% between reduced and original sets. The other measures were less robust, reaching 87.9% (mRECIST), 86.6% (Angle) and 79.9% (AUC), respectively. This suggests that it may be possible to reduce the number of replicates to 3 when studying drug response if necessary. However, we have seen that good estimates for the inter-replicate variability are important. This can be done better with 6 or more replicates and we therefore encourage the experimenters to continue PDX experiments with more replicates to maintain higher accuracy when possible.

### Clinical relevance of KuLGaP

We compared the cisplatin-vinorelbine combination treatment response in PDXs to data from 13 corresponding non-small cell lung carcinoma (NSCLC) patients receiving adjuvant platinum-based chemotherapy. For each of these patients, we considered both the time to recurrence and the growth curves of the corresponding PDX. The time to recurrence was measured from the time of starting adjuvant chemotherapy to either recurrence or last follow up.

We found that among patients whose corresponding PDX models were classified as responders by KuLGaP, the mean time to recurrence was 4.13 years, compared to 1.02 years in the group of non-responders according to KuLGaP. The difference (3.11 years) was the highest compared to all other methods (Table 1). We have found significant disagreement between the measures. There was unanimity between the responders in only four cases (three responders, one non-responder).

Due to the small sample size, it is difficult to assess statistical significance of our clinical validation, however the fact that there is a substantial difference in survival of the patients we predict as PDX responders, compared to other methods, is very encouraging in terms of clinical relevance of our measure compared to all other currently used approaches.

## Discussion

The problem of drug response prediction is incredibly important for the field of precision medicine, but is far from being solved and fraught with many obstacles. Patient derived xenografts are certainly a very appealing paradigm for drug response studies due to the ability to implant a patient’s tumor into a living organism (mouse) where it can potentially act as a realistic simulation of a given patient, and model the spectrum of clinical disease. Among their many applications, PDX can be used both for predicting response for individual patients through empiric drug treatment, and for identifying biomarker-response relationships across heterogenous collections representing the patient populations. In each use case, efficient testing of many individual PDX models and drugs, and accurate drug response quantification are of critical importance. In the former, false positive or negative predictions have a major impact, as patients have a limited opportunity for treatment, and avoidance of ineffective toxic therapy is crucial. In the latter, accurate response calls are necessary in order to identify or validate predictive biomarkers that can be used to guide patient selection or companion diagnostic development in clinical trials. As PDX are not quite the same as a patient and there is variation of response even among identical mice, it is essential to have a robust measure quantifying response from these experiments. Our work shows that none of the currently widely used response quantification measures take into account the full extent of the available experimental data, some ignore controls and others - variation among replicates. In this work we proposed a novel measure, KuLGaP, that provides a theoretically sound solution to this problem, which we have shown to be more selective on a large set of PDXs and concordant with patient outcomes in a small study.

Our exploration of real-world examples provides an insight into how we could improve other existing measures as well. For example, one way to make the mRECIST measure more selective would be to include stable disease (mSD) in the non-responder category. Unfortunately, this leads to false negative classifications: an extreme example of this is illustrated in Supplementary Figure 3.

The TGI measure is one of the widely used measures in the biomedical literature. Like the most commonly-used measures, TGI is computed based on the mean value of the replicates and then thresholded, especially in cases when the number of replicates is small, and therefore fails to take into account the variation between replicates. As discussed above, this can have a substantial impact on the resulting classification. Moreover, the TGI criterion only takes the first and last measurement into account and is therefore highly susceptible to measurement errors and fluctuations in the tumor size at the specific timepoints. One way to introduce at least some impact of the variance would be to calculate TGI individually for each control-treatment replicate pair and apply a suitable statistical test. However, this approach would not work well in models with relatively few replicates per model, since this would lead to a low power in the statistical testing. In order to reduce the impact of a measurement error at the end of the experiment, one could calculate the TGI criterion based on a few of the measurement points and then take a consensus measure. While it may result in an improvement, this solution will still suffer from not considering the variance across timepoints.

Overall, in our experience, there is no substitute for a measure that models all of the available data simultaneously, taking advantage of the multiple replicates for cases and controls; KuLGaP fulfills these criteria. We expect that introducing such a measure will lead to more faithful predictions of clinical outcomes, and biomarker-response relationships. We have thus created a simple to use web interface to assess the response for any PDX clinical experiments, kulgap.ca, that is equally easy to use for both clinicians, technicians and bio-statisticians, which we hope will result in wide uptake and reproducible results across drug response research.

## Online Methods

### Data preparation

At each measured time point of an experiment, we took tumor volume estimated from the tumor dimensions as the observed treatment response. The first day of drug administration was designated as the initial point of the experiment and we studied the growth curves from that point onwards. Next, the growth curve of each PDX replicate, in both treatment and control arms, was log-normalized to the tumor size at the starting day of the treatment. The treatment response was then assessed from these truncated log-normalized curves.

### KuLGaP

There are two steps to computing KuLGaP. In the first, two Gaussian Process (GP) models ^7^ are fitted, one for tumor treated PDX and one for controls. In the second step, we compute a symmetrised integrated version of the Kullback-Leibler (KL) divergence between the two GP models called Kullback Leibler (KL) divergence. KL is frequently used to compute the distance between two distributions. We assessed the significance of divergence between two models by computing KL divergences between all pairs of controls. Using this empirical distribution of divergences, we computed p-values of significance of response for each PDX model. Models with a p-value less than 0.05 were considered to have a statistically significant KL divergence were classified as responders.

#### Gaussian Processes

Recall that a set of random variables *X*_1_, …, *X*_*k*_ is said to be *jointly Gaussian* with mean vector μ ∈ ℝ^*k*^ and covariance matrix Σ ∈ ℝ^*k*×*k*^ if the joint density of *X*_1_, …, *X*_*k*_ is given by

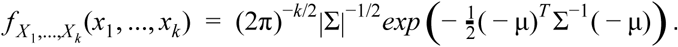

A *Gaussian Process* (*GP*) ^17^ on an interval [0, *T*] with *mean process m* : [0, *T*] → ℝ and *covariance kernel K* : [0, *T*] × [0, *T*] → ℝ can be considered as an infinite-dimensional analogue of the joint Gaussian distribution and is formally defined as a random function *X* : [0, *T*] → ℝ such that for any 0 < *t*_1_ < … < *t*_*k*_ < *T* the joint distribution of (*X*(*t*_1_), …, *X*(*t*_*k*_) is Gaussian with mean vector (*m*(*t*_1_), …, *m*(*t*_*k*_)) and covariance matrix Σ, where Σ_*i,j*_ = *K*(*t*_*i*_, *t*_*j*_) for all *i, j*.

Given a collection of measurements - such as tumor sizes measured for each replicate in a PDX experiment, separately for treatment and control, and a prior GP, one can use Bayes’ theorem to find the posterior distribution given the data, see also Bishop ^18^, Chapter 6.4. This was implemented using the GPy package ^19^ (http://github.com/SheffieldML/GPy). Due to its universality ^20^ and for theoretical reasons ^7^, the radial basis function (RBF) was chosen as the prior distribution, with a variance of 1 and a length scale of 10. This choice for a prior kernel leads to good fits of the data for the posterior distribution. Hyper parameter selection was performed by maximizing the likelihood, using the Broyden-Fletcher-Goldfarb-Shannon algorithm provided by the package, with seven restarts for each model. The schematic for our data analysis pipeline is given in Figure 1.

#### Kullback-Leibler Divergence and KuLGaP

The *Kullback-Leibler divergence* ^8^ *(also called relative entropy*) between two probability measures *P* and *Q* on a set is given by

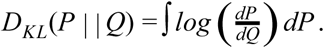

This is not symmetric, and it will be more convenient to work with the symmetrised version,

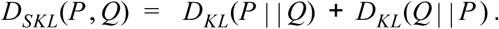

For two random processes, that is sequences of probability measures (μ _*t*_, ν_*t*_ : *t* ∈ [0, *T*]) indexed by a time interval, we define the integrated symmetrised KL divergence between them as:

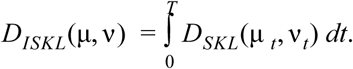

Consider now a particular PDX experiment with a given drug *D*, lasting a total of *T* days. We proceed as follows: First, fit a Gaussian process each to the treatment and the control replicates and denoting their distributions by μ^*T*^ = (μ_*t*_^*T*^ : *t* ∈ [0, *T*]) and μ^*C*^ = (μ_*t*_^*C*^ : *t* ∈ [0, *T*]) respectively, compute the integrated KL divergence *D*_*ISKL*_(μ^*T*^, μ^*C*^) between them. This quantity can be considered as a continuous estimate of the effect of drug *D*: the larger the KL divergence, the further away the treatment and control replicates are to one another, and therefore the larger an effect by drug *D*.

In order to test whether an observed KL value corresponds to a successful anticancer therapy we consider the null hypothesis *H*_0_ that the treatment and control GPs do not differ significantly and test it against the alternative hypothesis *H*_1_ that they do differ. We have chosen to estimate the distribution of a KL divergence under *H*_0_ empirically as follows. Since each control group does not receive any treatment, it is reasonable to assume that there is no effect. Therefore we have estimated the null distribution by computing empirical distribution by calculating the KL divergence between any pair of control groups from the NSCLC and colorectal PDX. This discrete distribution was then smoothed using a Gaussian kernel with bandwidth 0.27, which was selected via leave-one-out cross-validation by the statsmodels Python module ^21^. Finally, the KuLGaP measurement is calculated as the probability of obtaining a KL divergence value at least as large as the one obtained in the experiment (right tail probability/one-sided p-value). According to the empirical distribution we obtained, the critical values for the 0.1, 0.05 and 0.001 confidence levels were 5.61, 7.97 and 13.9, respectively. In particular, since we have chosen the 0.05 confidence level, an experiment was classified as a responder according to KuLGaP if and only if its KL divergence value was higher than 7.97. The observed values and our estimate of the probability distribution are illustrated in Supplementary Figure 7.

### Modified RECIST (mRECIST)

The Response Evaluation Criteria In Solid Tumors (RECIST) ^22^ is a framework of guidelines for evaluation of tumor response to anticancer therapies, based on linear dimensions of tumor lesions. Four classifications are possible, from the best to the worst outcome: complete response (CR), partial response (PR), stable disease (SD) and progressive disease (PD). The modified RECIST (mRECIST) ^1^ allows the classification based on tumor volume growth curves.

For each time *t*, we determined the relative volume change of the tumor with respect to its reference size *V* _0_, that is we calculated Δ*V* _*t*_ = (*V* _*t*_ − *V* _0_)/*V* _0_. The BestResponse is defined ^1^ to be the minimal value of Δ*V* _*t*_ for all times *t* after 3 days. Further, the running average of Δ*V* _0_, Δ*V* _1_, …, Δ*V* _*t*_ is calculated. The minimal value of this running average is called ^1^ BestAvgResponse. The quantities BestResponse and BestAvgResponse are then used to obtain the mRECIST classification, using the following thresholds ^1^:

- BestResponse < -95% and BestAvgResponse < -40%: *mCR* (modified complete response);
- BestResponse < -50% and BestAvgResponse < -20%: mPR (modified partial response);
- BestResponse < 35% and BestAvgResponse < 30%: mSD (modified stable disease);
- BestResponse > 35% or BestAvgResponse >30%: mPD (modified progressive disease).

Since the mRECIST criterion does not take into account the presence of multiple replicates, an mRECIST value is calculated for each replicate and a majority vote among replicate classifications is taken. Following Gao et al. ^1^, an mRECIST classification of mPD was considered as a non-responder, while all others as responders. It should be noted that by definition, mRECIST is not able to take into account the evolution over time (since it only considers the smallest observation), nor the variation between replicates.

### Area under the curve (AUC)

As done by Duan et al. ^2^, the area under the curve (AUC) under each replicate in the treatment and control groups was calculated. Then, p-values for group comparisons based on AUC were calculated using a one-tailed non-parametric Mann–Whitney test. A significance level of p<0.05 was used to classify each PDX model as either a responder (significant difference) or non-responder (no significant difference).

### Response angle

For each replicate in the treatment and control groups, the angle between the OLS best-fit of the normalized tumor curve and the line y=1 was calculated. Then, the same statistical test as described for the AUC was applied to compare pairwise mean angles of response ^2^, yielding a classification of each PDX model as either a responder (significant difference) or non-responder (no significant difference).

### The tumor growth inhibition (TGI)

The TGI is computed as follows: 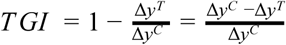, where Δ*y*^*C*^ and Δ*y*^*T*^ denote the mean difference between last and first measurement for the control and treatment groups respectively ^5^. Following established practice ^11–13^, we consider all PDX with a TGI value of more than 0.6 to be responders.

### Research Reproducibility

Our code and documentation are open-source and publicly available through the KulGaP GitHub repository (https://github.com/bhklab/pyKuLGaP). A detailed tutorial describing how to run our pipeline and reproduce our analysis results is available in the GitHub repository. A virtual machine reproducing the full software environment is available on Code Ocean ^23^. Our study complies with the guidelines outlined in ^24–26^. All the data are available in the form of XevaSet objects.

## Acknowledgements

JO was partially supported by a Horizon postdoctoral fellowship from Concordia University. The lung PDX resource has been supported by grants held by Tsao MS (Ontario Research Fund-Research Excellence Grant #03-020, Canadian Cancer Society Research Institute Grant #701595, and The Canadian Institutes of Health Research - Foundation Grant #148395). Pham NA and the Princess Margaret Living Biobank is funded by The Princess Margaret Cancer Foundation. AG, BHK, ET and JO were supported by a CCSRI and CHRP grant to AG and B.H.-K. DWC, B.H.-K. and ASM were supported by the Stand Up To Cancer Canada-Canadian Breast Cancer Foundation Breast Cancer Dream Team Research Funding, with supplemental support of the Ontario Institute for Cancer Research through funding provided by the Government of Ontario (Funding Award SU2C-AACR-DT-18-15). Stand Up To Cancer Canada is a program of the Entertainment Industry Foundation Canada. Research funding is administered by the American Association for Cancer Research International-Canada, the Scientific Partner of SU2C Canada. DWC and B.H.-K. were supported by The Terry Fox Research Institute, and B.H.-K. was supported by the Gattuso Slaight Personalized Cancer Medicine Fund at Princess Margaret Cancer Centre, the Canadian Institute of Health Research, and the Natural Sciences and Engineering Research Council. AG was supported by the Canadian Institute of Health Research, the Canadian Cancer Society, Terry Fox Foundation and the Natural Sciences and Engineering Research Council as well as CIFAR and the Amar Varma Family Chair in Biomedical Informatics and Artificial Intelligence.

## References

1. Gao, H. et al. High-throughput screening using patient-derived tumor xenografts to predict clinical trial drug response. Nat. Med. 21, 1318–1325 (2015).

2. Duan, F. et al. Area under the curve as a tool to measure kinetics of tumor growth in experimental animals. J. Immunol. Methods 382, 224–228 (2012).

3. Bertotti, A. et al. The genomic landscape of response to EGFR blockade in colorectal cancer. Nature 526, 263–267 (2015).

4. Yung-mae, M. Y. et al. Mouse PDX Trial Suggests Synergy of concurrent Inhibition of RAF and EGFR in Colorectal Cancer with BRAF or KRAS mutations. Clin. Cancer Res. clincanres–3250 (2017).

5. Guo, S., Jiang, X., Mao, B. & Li, Q.-X. The design, analysis and application of mouseclinical trials in oncology drug development. BMC Cancer 19, 718 (2019).

6. Laajala, T. D. et al. Optimized design and analysis of preclinical intervention studies in vivo. Sci. Rep. 6, 30723 (2016).

7. Williams, C. K. & Rasmussen, C. E. Gaussian processes for machine learning. the MIT Press 2, 4 (2006).

8. Kullback, S. & Leibler, R. A. On Information and Sufficiency. Ann. Math. Stat. 22, 79–86 (1951).

9. Pardo, M. C. & Vajda, I. About distances of discrete distributions satisfying the data processing theorem of information theory. IEEE Trans. Inf. Theory 43, 1288–1293 (1997).

10. Burnham, K. P. & Anderson, D. R. Model Selection and Multimodel Inference: A Practical Information-Theoretic Approach. (Springer Science & Business Media, 2003).

11. Hong, B. et al. Intra-tumour molecular heterogeneity of clear cell renal cell carcinoma reveals the diversity of the response to targeted therapies using patient-derived xenograft models. Oncotarget 8, 49839–49850 (2017).

12. Guo, S. et al. Cetuximab response in CRC patient-derived xenografts seems predicted by an expression based RAS pathway signature. Oncotarget 7, 50575–50581 (2016).

13. Guo, S., Mao, B. & Li, H. Q. Abstract 4534: Theory and methodology for the design and analysis of PDX mouse clinical trials. Cancer Res. 77, 4534–4534 (2017).

14. Stewart, E. L. et al. Clinical Utility of Patient-Derived Xenografts to Determine Biomarkers of Prognosis and Map Resistance Pathways in EGFR-Mutant Lung Adenocarcinoma. J. Clin. Oncol. 33, 2472–2480 (2015).

15. Brana, I. et al. Novel combinations of PI3K-mTOR inhibitors with dacomitinib or chemotherapy in PTEN-deficient patient-derived tumor xenografts. Oncotarget 8, 84659–84670 (2017).

16. Krepler, C. et al. A Comprehensive Patient-Derived Xenograft Collection Representing the Heterogeneity of Melanoma. Cell Rep. 21, 1953–1967 (2017).

17. Williams, C. K. I. & Rasmussen, C. E. Gaussian processes for machine learning. vol. 2 (MIT press Cambridge, MA, 2006).

18. Bishop, C. M. Pattern Recognition and Machine Learning. (Springer New York, 2016).

19. GPy, G. A gaussian process framework in python. (2012).

20. Micchelli, C. A., Xu, Y. & Zhang, H. On translation invariant operators which preserve the B-spline recurrence. Adv. Comput. Math. 28, 157–169 (2008).

21. Seabold, S. & Perktold, J. Statsmodels: Econometric and statistical modeling with python. in Proceedings of the 9th Python in Science Conference vol. 57 61 (Scipy, 2010).

22. Therasse, P. et al. New guidelines to evaluate the response to treatment in solid tumors. European Organization for Research and Treatment of Cancer, National Cancer Institute of the United States, National Cancer Institute of Canada. J. Natl. Cancer Inst. 92, 205–216 (2000).

23. Haibe-Kains, B. et al. KuLGaP: A Selective Measure for Assessing Therapy Response in Patient-Derived Xenografts. (Code Ocean, 2020). doi: 10.24433/CO.8958866.V1.

24. Sandve, G. K., Nekrutenko, A., Taylor, J. & Hovig, E. Ten simple rules for reproducible computational research. PLoS Comput. Biol. 9, e1003285 (2013).

25. Gentleman, R. Reproducible research: a bioinformatics case study. Stat. Appl. Genet. Mol. Biol. 4, Article2 (2005).

26. Stroup, D. F. et al. Meta-analysis of Observational Studies in Epidemiology: A Proposal for Reporting. JAMA 283, 2008–2012 (2000).

