## Supplementary Tables and Figures for "KuLGaP: A Selective Measure for Assessing Therapy Response in Patient-Derived Xenografts"

Janosch Ortmann[1,13], Ladislav Rampášek[2,6,7], Elijah Tai[2], Arvind Singh Mer[3,4], Ruoshi Shi[3], Erin L Stewart[3], Celine Mascaux[3], Aline Fares[3], Nhu-An Pham[3], Gangesh Beri[3], Christopher Eeles[3], Denis Tkachuk[3], Chantal Ho[3], Shingo Sakashita[3], Jessica Weiss[3], Xiaoqian Jiang[5], Geoffrey Liu[3], David W. Cescon[3], Catherine O'Brien[3,4,9,10,11], Sheng Guo[5], Ming-Sound Tsao[3], Benjamin Haibe-Kains[2,3,4,6,8], Anna Goldenberg[2,6,7,12]

[1] Département AOTI, Université du Québec à Montréal, Montreal, QC, H2X3X2, Canada
[2] Department of Computer Science, University of Toronto, Toronto, ON, M5S2E4, Canada
[3] Princess Margaret Cancer Centre, University Health Network, Toronto, ON, M5G1L7, Canada
[4] Department of Medical Biophysics, University of Toronto, Toronto, ON, M5G1L7, Canada
[5] Crown Bioscience Taicang, Inc., Taicang, Jiangsu, China
[6] Vector Institute for Artificial Intelligence, Toronto, ON, M5G1M1, Canada
[7] Hospital for Sick Children, Toronto, ON, M5G1X8, Canada
[8] Ontario Institute for Cancer Research, Toronto, ON, M5G1L7, Canada
[9] Department of Laboratory Medicine and Pathobiology, University of Toronto, Toronto, ON M5S1A8, Canada
[10] Department of Physiology, University of Toronto, Toronto, ON, M5G1L7, Canada
[11] Department of Surgery, Toronto General Hospital, Toronto, ON, M5G2C4, Canada
[12] CIFAR, Toronto, ON, M5G1M1, Canada
[13] Group for research in decision analysis (GERAD), Université de Montréal, Montreal, QC, H3T1J4, Canada

| Number of experiments | Type of cancer | Description | Drugs tested |
|---|---|---|---|
| 116 | Non-small cell lung carcinoma (NSCLC) | Retrospectively collected data from the Princess Margaret Living Biobank - PDX core (University Health Network) . 1260 individual mouse models from 30 patients and 3 cell lines. On average, 7 PDX replicates per model. The mice were implanted with about 64mm$^3$ tumour material. Once the tumor volume reached 150mm$^3$ to 200mm$^3$ the mice were randomized to treatment or control group. Tumor volume was measured every 3rd day until 90 days had passed. If tumor volume reached more than 2000mm$^3$, the experiment was terminated. | In total, 36 different drugs and drug combinations were tested. On average, 3 different drugs were tested on each PDX sample |
| 13 | colorectal | | PDXs were treated with evofosfamide (Evo), a hypoxia-activated prodrug (HAP), 5-Fluorouracil (5-FU) or chemoradiotherapy (5-FU and radiation; CRT). |
| 200 | various | Data from Crown Bioscience Inc. Mice were inoculated when a tumor was between 100 to 300mm$^3$. Tumor volume was measured twice every week until it reached 3000mm$^3$ or doubled or the experiment ran for 60 days. For each treatment group, 3 to 10 replicates were used. | various |

**Supplementary Table 1**. Summary of PDX model collections used in this paper.

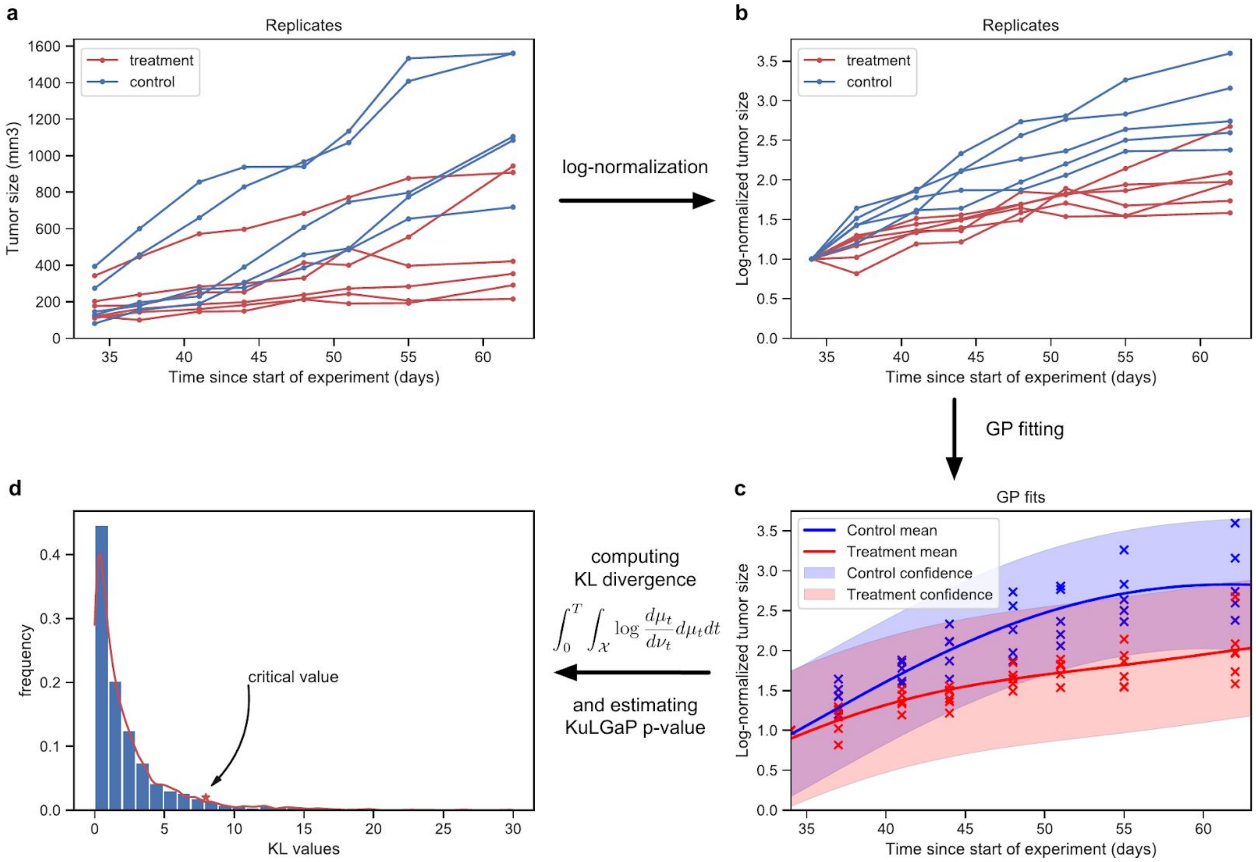


**Supplementary Figure 1:** ***The KuLGaP pipeline***. **a**,**b**, Tumor volumes of each replicate are normalized to the volume at the starting day of the treatment (here it is day 34) and log-transformed. **c**, A Gaussian process is fitted to the treatment and the control group of replicates. **d**, KL divergence between the two GPs serves as a numerical estimator for how different the treatment and control groups are. We arrive at the KuLGaP by computing a *p*-value against the estimated null distribution of KL values.

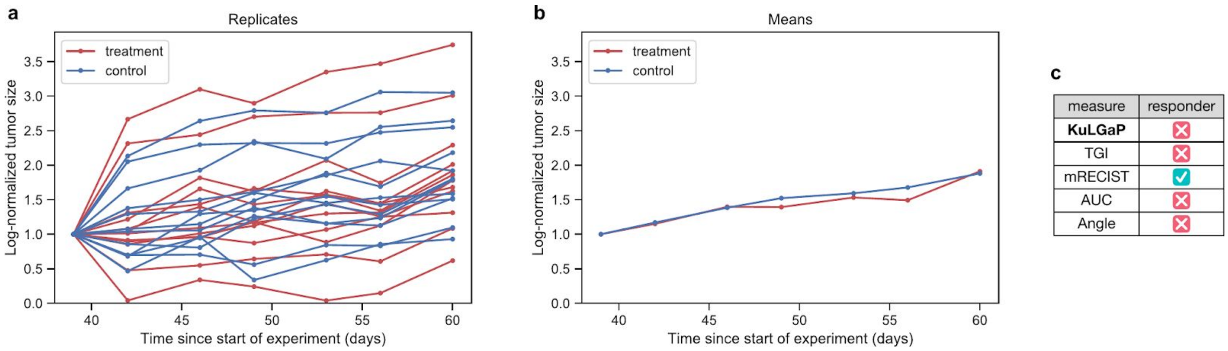


**Supplementary Figure 2:** ***Optimistic classification by mRECIST.*** Log-normalised tumor-control curves for a breast cancer PDX model[1] with 12 replicates of the same model treated with gedatolisib. On the left panel, all replicates are shown and on the right panel, the pointwise means across all replicates for control and treatment groups respectively. All considered measures, except for mRECIST, classify this experiment as a non-responder. (**p36)**

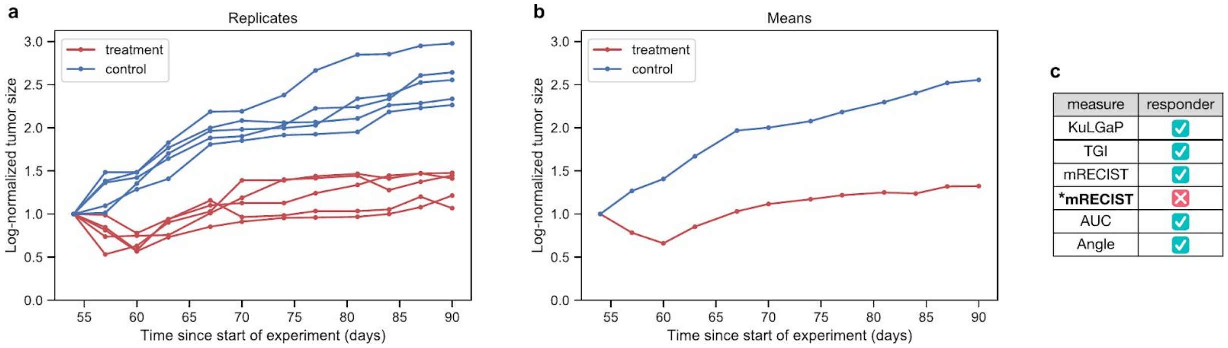


**Supplementary Figure 3:** ***Downside of altering mRECIST classification to be more conservative***. An experiment with five replicates treated with BKM-120 and five control replicates; shown in logarithmic scale. KuLGaP and all four baseline measures agree on a "responder" classification. However, altering mRECIST classification to be more conservative by considering only mPR and mCR as a response (*mRECIST) leads to a false negative, demonstrating a clear loss of sensitivity. (**p62**)

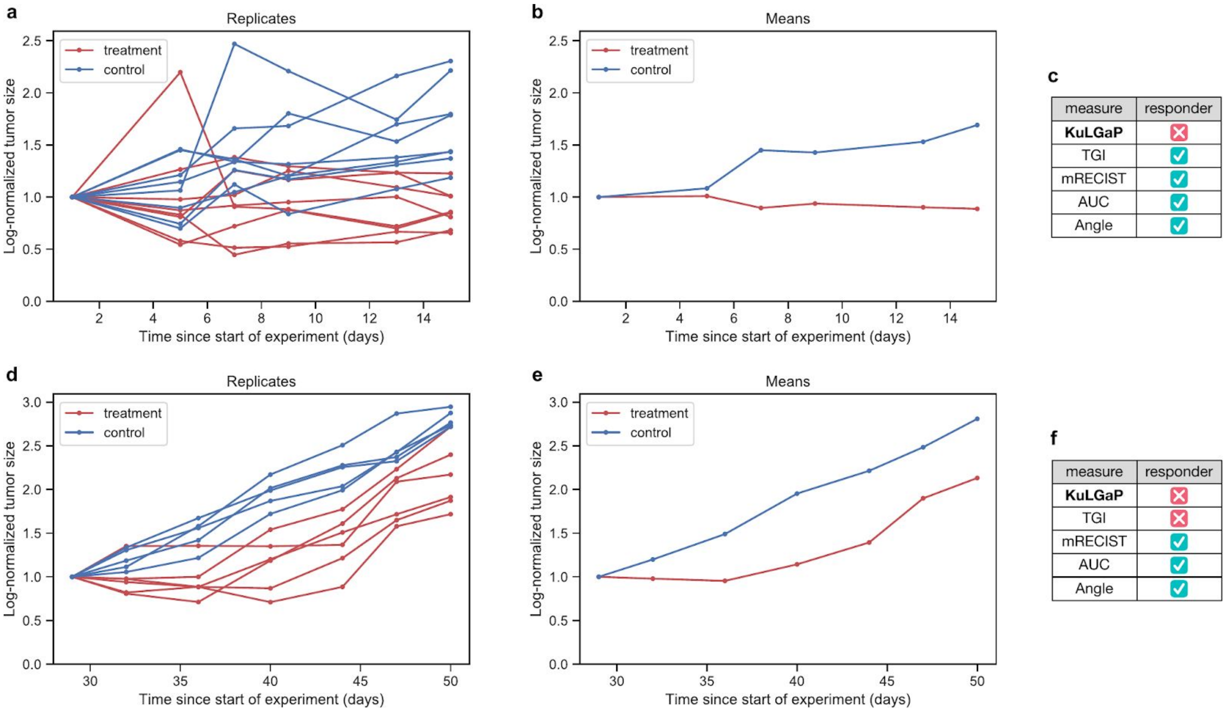


**Supplementary Figure 4:** ***Importance of considering the control group.*** **a**,**b**,**c**, Colorectal PDX model with eight treatment replicates, treated with evofosfamide and eight control replicates; shown in logarithmic scale. All measures apart from KuLGaP agree on a responder classification. **d**,**e**,**f**, NSCLC PDX model with six treatment replicates, treated with BKM-120, and five control replicates; shown in logarithmic scale. All measures except KuLGaP and TGI agree on a responder classification.

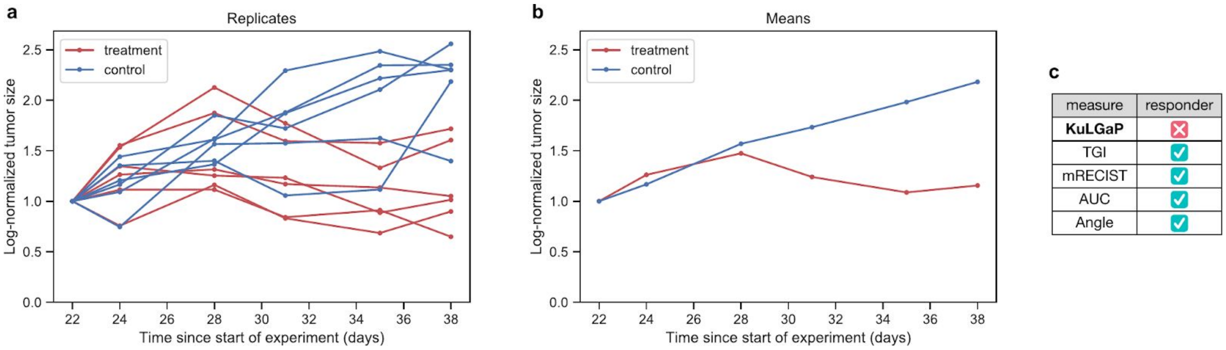


**Supplementary Figure 5:** ***Importance of accounting for variance.*** **a**, Log-normalized growth curves of a NSCLC PDX model with 6 replicates treated with vandetanib (treatment replicates in red, controls in blue). **b**, Pointwise means across replicates for control and treatment groups respectively. **c**, Computed response classifications by all compared response measures.

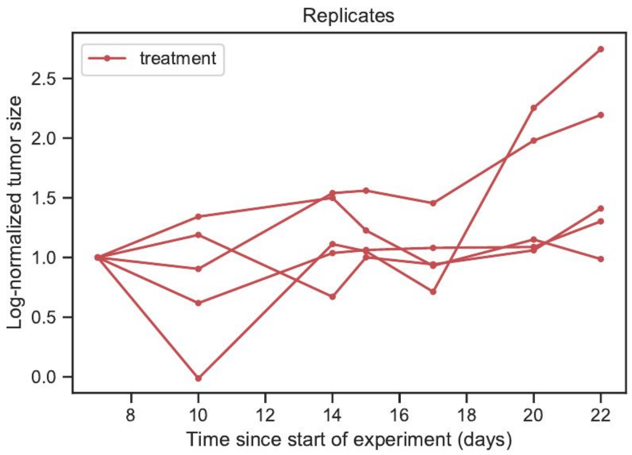


**Supplementary Figure 6:** ***Variety of mRECIST classifications in the replicates***. Log-normalized tumor growth curves for five replicates treated with erlotinib. Their difference after the first measurement (day 10) translates into three different mRECIST classification results: one experiment is classified as mSD, two as mPR, and two as mCR. (***p51***)

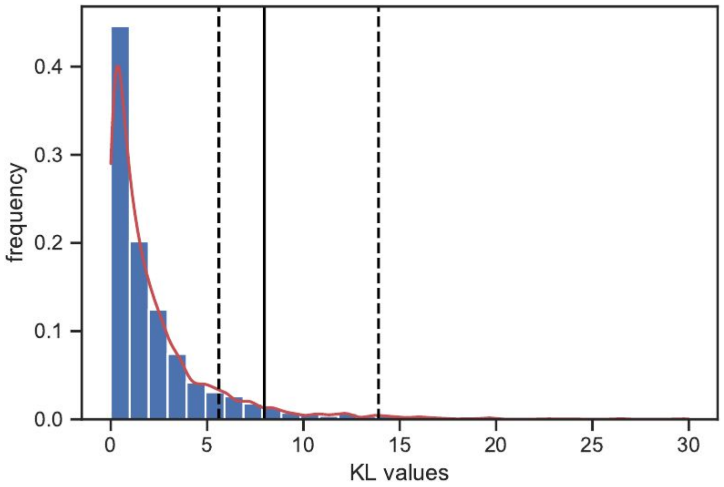


**Supplementary Figure 7:** ***The null distribution of KL values used in KuLGaP.*** Histogram of observed Kullback-Leibler values between controls (blue) and our estimation of the probability distribution (red). The solid vertical line at 7.97 indicates the KuLGaP's threshold for significance at the 0.05 level, while the dashed lines at 5.61 and 13.9 indicate the 0.1 and 0.001 thresholds, respectively.

| ID | KuLGaP | mRECIST-Novartis | Angle | AUC | TGI |
|---|---|---|---|---|---|
| 1 | 1 | 1 | 1 | 1 | -1 |
| 2 | -1 | 1 | 1 | 1 | -1 |
| 3 | -1 | 1 | 1 | 1 | -1 |
| 4 | 1 | 1 | 1 | 1 | 1 |
| 5 | -1 | 1 | 1 | 1 | 1 |
| 6 | -1 | 1 | 1 | 1 | 1 |
| 7 | -1 | -1 | -1 | -1 | -1 |
| 8 | -1 | -1 | -1 | 1 | -1 |
| 9 | -1 | -1 | 1 | 1 | 1 |
| 10 | -1 | -1 | -1 | 1 | -1 |
| 11 | -1 | 0 | 1 | 1 | 0 |
| 12 | -1 | -1 | -1 | -1 | -1 |
| 13 | -1 | -1 | 1 | 1 | 1 |
| 14 | -1 | -1 | 1 | 1 | -1 |
| 15 | -1 | -1 | -1 | -1 | -1 |
| 16 | -1 | -1 | -1 | -1 | -1 |
| 17 | -1 | 1 | 1 | 1 | -1 |
| 18 | -1 | -1 | 1 | -1 | -1 |
| 19 | -1 | 1 | 1 | -1 | 1 |
| 20 | -1 | 1 | 1 | 1 | 1 |
| 21 | -1 | 1 | -1 | -1 | -1 |
| 22 | -1 | 1 | -1 | -1 | -1 |
| 23 | -1 | 1 | 1 | 1 | 1 |
| 24 | -1 | 1 | -1 | -1 | -1 |
| 25 | -1 | 1 | 1 | 1 | -1 |
| 26 | -1 | -1 | -1 | 1 | -1 |
| 27 | -1 | 1 | -1 | -1 | -1 |
| 28 | -1 | 1 | 1 | 1 | -1 |
| 29 | -1 | 1 | 1 | 1 | -1 |
| 30 | -1 | 1 | 1 | 1 | -1 |
| 31 | -1 | 0 | 1 | 1 | 0 |
| 32 | -1 | 1 | 1 | -1 | -1 |
| 33 | -1 | 1 | 1 | -1 | -1 |

| | | | | | |
|---|---|---|---|---|---|
| 34 | -1 | 1 | 1 | 1 | -1 |
| 35 | -1 | 1 | -1 | -1 | -1 |
| 36 | -1 | -1 | -1 | -1 | -1 |
| 37 | -1 | 1 | -1 | -1 | -1 |
| 38 | -1 | -1 | 1 | -1 | -1 |
| 39 | -1 | -1 | 1 | 1 | -1 |
| 40 | -1 | -1 | -1 | -1 | -1 |
| 41 | -1 | -1 | -1 | -1 | -1 |
| 42 | -1 | -1 | -1 | 1 | -1 |
| 43 | -1 | -1 | -1 | -1 | -1 |
| 44 | -1 | -1 | -1 | -1 | -1 |
| 45 | -1 | 1 | 1 | 1 | -1 |
| 46 | -1 | -1 | 1 | 1 | 1 |
| 47 | 1 | 1 | 1 | 1 | 1 |
| 48 | -1 | 1 | 1 | 1 | 1 |
| 49 | -1 | -1 | 1 | 1 | -1 |
| 50 | -1 | 1 | 1 | -1 | -1 |
| 51 | -1 | -1 | 1 | 1 | 1 |
| 52 | -1 | 1 | -1 | 1 | -1 |
| 53 | -1 | -1 | -1 | -1 | -1 |
| 54 | -1 | 1 | 1 | 1 | -1 |
| 55 | -1 | 1 | 1 | -1 | -1 |
| 56 | 1 | 1 | 1 | 1 | 1 |
| 57 | 1 | 1 | 1 | 1 | 1 |
| 58 | -1 | 1 | 1 | -1 | -1 |
| 59 | -1 | 1 | -1 | -1 | -1 |
| 60 | -1 | 1 | -1 | -1 | -1 |
| 61 | -1 | 1 | 1 | 1 | 1 |
| 62 | 1 | 1 | 1 | 1 | 1 |
| 63 | 1 | 1 | 1 | 1 | -1 |
| 64 | 1 | 1 | 1 | 1 | -1 |
| 65 | 1 | 1 | 1 | 1 | -1 |
| 66 | 1 | 1 | 1 | 1 | 1 |
| 67 | 1 | 1 | 1 | 1 | 1 |

| | | | | | |
|---|---|---|---|---|---|
| 68 | -1 | 1 | -1 | -1 | -1 |
| 69 | -1 | -1 | -1 | -1 | -1 |
| 70 | 1 | 1 | 1 | 1 | 1 |
| 71 | 1 | 1 | 1 | 1 | 1 |
| 72 | -1 | 1 | 1 | -1 | 1 |
| 73 | -1 | 1 | -1 | -1 | -1 |
| 74 | 1 | 1 | -1 | -1 | -1 |
| 75 | -1 | 1 | -1 | -1 | -1 |
| 76 | -1 | 1 | -1 | -1 | -1 |
| 77 | -1 | 1 | 1 | -1 | -1 |
| 78 | 1 | 1 | 1 | 1 | 1 |
| 79 | 1 | 1 | 1 | 1 | 1 |
| 80 | -1 | 1 | 1 | 0 | -1 |
| 81 | -1 | 1 | 1 | 0 | 1 |
| 82 | -1 | 1 | 1 | 0 | -1 |
| 83 | -1 | 1 | 1 | 0 | 1 |
| 84 | -1 | 1 | 1 | 0 | -1 |
| 85 | -1 | 1 | 1 | 0 | 1 |
| 86 | -1 | 1 | 1 | 0 | 1 |
| 87 | 1 | 1 | 1 | 1 | 1 |
| 88 | -1 | 1 | -1 | 1 | 1 |
| 89 | -1 | -1 | -1 | -1 | -1 |
| 90 | -1 | 1 | 1 | -1 | -1 |
| 91 | -1 | 1 | 1 | 0 | -1 |
| 92 | -1 | 1 | 1 | 1 | -1 |
| 93 | -1 | -1 | 1 | -1 | -1 |
| 94 | -1 | 1 | 1 | -1 | -1 |
| 95 | -1 | 1 | 1 | -1 | -1 |
| 96 | -1 | 1 | 1 | -1 | -1 |
| 97 | -1 | -1 | 1 | 0 | -1 |
| 98 | 1 | 1 | 1 | 0 | -1 |
| 99 | -1 | 1 | 1 | 1 | -1 |
| 100 | -1 | 1 | -1 | -1 | -1 |
| 101 | 1 | 1 | 1 | 1 | 1 |

| | | | | | |
|---|---|---|---|---|---|
| 102 | -1 | 1 | 1 | 1 | -1 |
| 103 | -1 | 1 | 1 | 1 | -1 |
| 104 | -1 | -1 | 1 | 1 | -1 |
| 105 | -1 | 1 | -1 | 1 | 1 |
| 106 | 1 | -1 | 1 | 1 | 1 |
| 107 | -1 | -1 | -1 | 1 | -1 |
| 108 | -1 | -1 | 1 | -1 | -1 |
| 109 | -1 | 1 | -1 | 1 | -1 |
| 110 | -1 | -1 | -1 | -1 | -1 |
| 111 | -1 | 1 | -1 | -1 | -1 |
| 112 | -1 | -1 | -1 | 1 | -1 |
| 113 | 1 | 1 | 1 | 1 | 1 |
| 114 | 1 | 1 | 1 | 1 | 1 |
| 115 | 1 | 1 | 1 | 1 | 1 |
| 116 | 1 | 1 | 1 | 1 | 1 |
| 117 | -1 | 1 | 1 | 1 | -1 |
| 118 | -1 | 1 | -1 | -1 | -1 |
| 119 | -1 | 1 | 1 | 1 | 1 |
| 120 | -1 | 1 | 1 | 1 | -1 |
| 121 | -1 | 1 | 1 | 1 | -1 |
| 122 | -1 | 1 | 1 | 1 | -1 |
| 123 | -1 | -1 | 1 | -1 | -1 |
| 124 | -1 | -1 | 1 | 1 | -1 |
| 125 | -1 | 1 | -1 | 1 | -1 |
| 126 | -1 | 1 | 1 | 1 | -1 |
| 127 | -1 | 1 | 1 | 1 | 1 |
| 128 | -1 | -1 | 1 | 1 | -1 |
| 129 | -1 | 1 | 1 | 1 | 1 |
| 130 | -1 | 1 | 1 | 1 | 1 |
| 131 | -1 | 1 | 1 | 1 | 1 |
| 132 | -1 | 1 | 1 | 1 | 1 |
| 133 | 1 | 1 | 1 | 1 | 1 |
| 134 | 1 | 1 | 1 | 1 | 1 |
| 135 | 1 | 1 | 1 | 1 | 1 |

| | | | | | |
|---|---|---|---|---|---|
| 136 | 1 | 1 | 1 | 1 | 1 |
| 137 | -1 | 1 | 1 | 1 | 1 |
| 138 | 1 | 1 | 1 | 1 | 1 |
| 139 | -1 | 1 | 1 | 1 | 1 |
| 140 | -1 | 1 | 1 | -1 | 1 |
| 141 | -1 | 1 | 1 | 1 | 1 |
| 142 | -1 | 1 | 1 | 1 | 1 |
| 143 | -1 | 1 | 1 | 1 | 1 |
| 144 | -1 | 1 | 1 | 1 | 1 |
| 145 | -1 | 1 | 1 | 1 | 1 |
| 146 | 1 | 1 | 1 | 1 | 1 |
| 147 | 1 | 1 | 1 | 1 | 1 |
| 148 | -1 | 1 | 1 | 1 | 1 |
| 149 | -1 | 1 | 1 | 1 | 1 |
| 150 | -1 | 1 | 1 | -1 | 1 |
| 151 | 1 | 1 | 1 | 1 | 1 |
| 152 | -1 | 1 | 1 | 1 | 1 |
| 153 | 1 | 1 | 1 | 1 | 1 |
| 154 | 1 | 1 | 1 | 1 | 1 |
| 155 | -1 | 1 | 1 | 1 | 1 |
| 156 | 1 | 1 | 1 | 1 | 1 |
| 157 | -1 | 1 | 1 | 1 | 1 |
| 158 | 1 | 1 | 1 | 1 | 1 |
| 159 | 1 | 1 | 1 | 1 | 1 |
| 160 | -1 | 1 | 1 | 1 | 1 |
| 161 | -1 | -1 | 1 | 1 | 1 |
| 162 | -1 | -1 | 1 | 1 | 1 |
| 163 | 1 | -1 | 1 | 1 | 1 |
| 164 | -1 | 1 | 1 | 1 | 1 |
| 165 | -1 | -1 | 1 | 1 | 1 |
| 166 | 1 | -1 | 1 | 1 | 1 |
| 167 | -1 | -1 | 1 | 1 | 1 |
| 168 | -1 | 1 | 1 | 1 | 1 |
| 169 | -1 | 1 | 1 | 1 | 1 |

| | | | | | |
|---|---|---|---|---|---|
| 170 | 1 | 1 | 1 | 1 | 1 |
| 171 | -1 | -1 | 1 | 1 | 1 |
| 172 | -1 | -1 | 1 | 1 | 1 |
| 173 | 1 | 1 | 1 | 1 | 1 |
| 174 | -1 | -1 | 1 | 1 | 1 |
| 175 | -1 | 1 | 1 | -1 | 1 |
| 176 | 1 | 1 | 1 | 1 | 1 |
| 177 | -1 | 1 | 1 | 1 | 1 |
| 178 | -1 | 1 | 1 | 1 | 1 |
| 179 | -1 | 1 | 1 | 1 | 1 |
| 180 | 1 | 1 | -1 | -1 | 1 |
| 181 | -1 | 1 | -1 | -1 | 1 |
| 182 | 1 | 1 | 1 | 1 | 1 |
| 183 | -1 | -1 | -1 | -1 | 1 |
| 184 | -1 | 1 | -1 | -1 | 1 |
| 185 | -1 | 1 | 1 | 1 | 1 |
| 186 | -1 | -1 | 1 | 1 | 1 |
| 187 | -1 | -1 | 1 | 1 | 1 |
| 188 | -1 | 1 | 1 | 1 | 1 |
| 189 | 1 | 1 | 1 | 1 | 1 |
| 190 | -1 | 1 | -1 | -1 | 1 |
| 191 | -1 | -1 | 1 | 1 | 1 |
| 192 | -1 | -1 | 1 | 1 | 1 |
| 193 | -1 | 1 | 1 | 1 | 1 |
| 194 | -1 | -1 | 1 | 1 | 1 |
| 195 | -1 | 1 | 1 | -1 | 1 |
| 196 | -1 | 1 | 1 | 1 | 1 |
| 197 | -1 | -1 | -1 | -1 | 1 |
| 198 | -1 | 1 | 1 | 1 | 1 |
| 199 | -1 | -1 | 1 | 1 | 1 |
| 200 | -1 | -1 | 1 | 1 | 1 |
| 201 | -1 | 1 | 1 | 1 | 1 |
| 202 | -1 | 1 | 1 | 1 | 1 |
| 203 | -1 | -1 | 1 | 1 | 1 |

| | | | | | |
|---|---|---|---|---|---|
| 204 | -1 | 1 | 1 | 1 | 1 |
| 205 | -1 | 1 | 1 | 1 | 1 |
| 206 | -1 | 1 | 1 | 1 | 1 |
| 207 | 1 | 1 | 1 | 1 | 1 |
| 208 | -1 | -1 | 1 | 1 | 1 |
| 209 | -1 | 1 | 1 | 1 | 1 |
| 210 | 1 | -1 | 1 | 1 | 1 |
| 211 | -1 | -1 | 1 | 1 | 1 |
| 212 | -1 | 1 | 1 | 1 | 1 |
| 213 | 1 | 1 | 1 | 1 | 1 |
| 214 | -1 | -1 | 1 | 1 | 1 |
| 215 | -1 | 1 | -1 | 1 | 1 |
| 216 | -1 | -1 | 1 | -1 | 1 |
| 217 | -1 | -1 | 1 | 1 | 1 |
| 218 | -1 | 1 | 1 | 1 | 1 |
| 219 | -1 | 1 | 1 | 1 | 1 |
| 220 | -1 | 1 | 1 | 1 | 1 |
| 221 | -1 | 1 | 1 | -1 | 1 |
| 222 | -1 | -1 | 1 | 1 | 1 |
| 223 | -1 | 1 | -1 | -1 | 1 |
| 224 | -1 | -1 | 1 | 1 | -1 |
| 225 | -1 | 1 | 1 | 1 | -1 |
| 226 | -1 | 1 | 1 | 1 | -1 |
| 227 | -1 | 1 | -1 | -1 | -1 |
| 228 | -1 | -1 | 1 | 1 | -1 |
| 229 | -1 | 1 | 1 | -1 | -1 |
| 230 | -1 | -1 | 1 | -1 | -1 |
| 231 | -1 | -1 | -1 | 1 | -1 |
| 232 | -1 | 1 | 1 | 1 | -1 |
| 233 | -1 | -1 | 1 | 1 | -1 |
| 234 | -1 | 1 | 1 | 1 | -1 |
| 235 | -1 | -1 | -1 | -1 | -1 |
| 236 | -1 | -1 | 1 | -1 | -1 |
| 237 | -1 | -1 | -1 | -1 | -1 |

| | | | | | |
|---|---|---|---|---|---|
| 238 | -1 | 1 | 1 | 1 | -1 |
| 239 | -1 | 1 | 1 | 1 | -1 |
| 240 | -1 | -1 | 1 | -1 | -1 |
| 241 | -1 | 1 | 1 | 1 | -1 |
| 242 | -1 | 1 | 1 | 1 | -1 |
| 243 | -1 | -1 | 1 | 1 | -1 |
| 244 | -1 | -1 | 1 | 1 | -1 |
| 245 | -1 | -1 | 1 | 1 | -1 |
| 246 | -1 | -1 | 1 | 1 | -1 |
| 247 | -1 | -1 | -1 | -1 | -1 |
| 248 | -1 | -1 | -1 | -1 | -1 |
| 249 | -1 | -1 | -1 | -1 | -1 |
| 250 | -1 | -1 | 1 | -1 | -1 |
| 251 | -1 | -1 | -1 | -1 | -1 |
| 252 | -1 | -1 | -1 | -1 | -1 |
| 253 | -1 | 1 | -1 | -1 | -1 |
| 254 | -1 | -1 | 1 | 1 | -1 |
| 255 | -1 | 1 | -1 | -1 | -1 |
| 256 | -1 | -1 | -1 | -1 | -1 |
| 257 | -1 | -1 | -1 | -1 | -1 |
| 258 | -1 | -1 | -1 | 1 | -1 |
| 259 | -1 | -1 | 1 | 1 | -1 |
| 260 | -1 | -1 | -1 | -1 | -1 |
| 261 | -1 | -1 | 1 | -1 | -1 |
| 262 | -1 | -1 | -1 | 1 | -1 |
| 263 | -1 | 1 | 1 | 1 | -1 |
| 264 | -1 | -1 | 1 | 1 | -1 |
| 265 | -1 | 1 | -1 | -1 | -1 |
| 266 | -1 | -1 | -1 | -1 | -1 |
| 267 | -1 | -1 | 1 | -1 | -1 |
| 268 | -1 | -1 | -1 | -1 | -1 |
| 269 | -1 | -1 | -1 | 1 | -1 |
| 270 | -1 | -1 | -1 | -1 | -1 |
| 271 | -1 | -1 | -1 | -1 | -1 |

| | | | | | |
|---|---|---|---|---|---|
| 272 | -1 | -1 | -1 | -1 | -1 |
| 273 | -1 | 1 | -1 | -1 | -1 |
| 274 | -1 | -1 | -1 | -1 | -1 |
| 275 | -1 | -1 | -1 | 1 | -1 |
| 276 | -1 | -1 | -1 | -1 | -1 |
| 277 | -1 | -1 | -1 | -1 | -1 |
| 278 | -1 | -1 | -1 | -1 | -1 |
| 279 | -1 | -1 | -1 | -1 | -1 |
| 280 | -1 | 1 | -1 | -1 | -1 |
| 281 | -1 | 1 | -1 | -1 | -1 |
| 282 | -1 | -1 | -1 | -1 | -1 |
| 283 | -1 | -1 | 1 | 1 | -1 |
| 284 | -1 | -1 | -1 | -1 | -1 |
| 285 | -1 | 1 | 1 | -1 | -1 |
| 286 | -1 | -1 | -1 | -1 | -1 |
| 287 | -1 | -1 | -1 | -1 | -1 |
| 288 | -1 | 1 | 1 | -1 | -1 |
| 289 | -1 | -1 | -1 | -1 | -1 |
| 290 | -1 | -1 | -1 | 1 | -1 |
| 291 | -1 | -1 | -1 | -1 | -1 |
| 292 | -1 | 1 | 1 | -1 | -1 |
| 293 | -1 | -1 | -1 | -1 | -1 |
| 294 | -1 | 1 | -1 | -1 | -1 |
| 295 | -1 | -1 | -1 | -1 | -1 |
| 296 | -1 | -1 | -1 | -1 | -1 |
| 297 | -1 | -1 | -1 | -1 | -1 |
| 298 | -1 | 1 | -1 | -1 | -1 |
| 299 | -1 | -1 | -1 | -1 | -1 |
| 300 | -1 | -1 | -1 | -1 | -1 |
| 301 | -1 | -1 | -1 | -1 | -1 |
| 302 | -1 | -1 | -1 | -1 | -1 |
| 303 | -1 | -1 | -1 | -1 | -1 |
| 304 | -1 | -1 | -1 | -1 | -1 |
| 305 | -1 | 1 | -1 | -1 | -1 |

| | | | | | |
|---|---|---|---|---|---|
| 306 | -1 | -1 | -1 | -1 | -1 |
| 307 | -1 | -1 | -1 | -1 | -1 |
| 308 | -1 | -1 | -1 | -1 | -1 |
| 309 | -1 | -1 | -1 | -1 | -1 |
| 310 | -1 | -1 | -1 | -1 | -1 |
| 311 | -1 | -1 | -1 | -1 | -1 |
| 312 | -1 | -1 | -1 | -1 | -1 |
| 313 | -1 | -1 | -1 | -1 | -1 |
| 314 | -1 | 1 | -1 | -1 | -1 |
| 315 | -1 | -1 | -1 | 1 | -1 |
| 316 | -1 | -1 | -1 | -1 | -1 |
| 317 | -1 | -1 | 1 | -1 | -1 |
| 318 | -1 | 1 | -1 | -1 | -1 |
| 319 | -1 | 1 | -1 | -1 | -1 |
| 320 | -1 | -1 | -1 | -1 | -1 |
| 321 | -1 | 1 | 1 | 1 | -1 |
| 322 | -1 | -1 | 1 | 1 | -1 |
| 323 | -1 | -1 | -1 | -1 | -1 |
| 324 | -1 | -1 | -1 | -1 | -1 |
| 325 | -1 | -1 | 1 | 1 | -1 |
| 326 | -1 | -1 | -1 | -1 | -1 |
| 327 | -1 | -1 | -1 | -1 | -1 |
| 328 | -1 | -1 | 1 | 1 | -1 |
| 329 | -1 | 1 | 1 | 1 | -1 |

**Supplementary Table 2**. Classification of all PDX models. Legend: -1 is a non-responder; 1 is a responder; 0 means the measure failed to run.

### References

1. Brana, I. *et al.* Novel combinations of PI3K-mTOR inhibitors with dacomitinib or chemotherapy in PTEN-deficient patient-derived tumor xenografts. *Oncotarget* **8**, 84659–84670 (2017).
